# Delayed recovery and host specialization may spell disaster for coral-fish mutualism

**DOI:** 10.1101/2022.06.01.494455

**Authors:** Catheline Y.M. Froehlich, O. Selma Klanten, Martin L. Hing, Mark Dowton, Marian Y.L. Wong

## Abstract

Mutualisms are prevalent in many ecosystems, yet little is known about how symbioses are affected by multiple disturbances. Here we show delayed recovery for 13 coral-dwelling goby fishes (genus *Gobiodon*) compared with their host *Acropora* corals following 4 consecutive cyclones and heatwaves. While corals became twice as abundant 3 years post-disturbances, their symbiotic gobies were only half as abundant relative to pre-disturbances and half of the goby species disappeared. Although goby species preferred particular coral species, surviving goby species shifted hosts to newly abundant coral species when their preferred hosts became rare. As host specialization is key for goby fitness, shifting hosts may have negative fitness consequences for gobies and corals alike and affect their survival in response to environmental changes. Our study demonstrates that mutualist partners do not respond identically to multiple disturbances, and that goby host plasticity, while potentially detrimental, may be the only possibility for early recovery.

## Introduction

In the face of climate change, multiple consecutive disturbances are becoming increasingly prevalent globally, and ecosystem stability is being threatened as a result^1,2^. Relationships between organisms are important for maintaining ecosystem balance and diversity during these challenging times, especially when one of these organisms is a habitat-forming foundation species, e.g. conifers, kelps, and corals^3,4^. Mutually beneficial symbioses (here termed ‘mutualisms’) often promote the survival of foundation and partner species, but anthropogenic disturbances are adding extreme pressures on these relationships^4,5^. A key question to arise is: will organisms in mutualisms respond similarly to consecutive disturbances, and what factors are important in the persistence of both partners^6^?

For symbioses in which one organism relies on the other for limiting resources like food and shelter, the host species is a key determinant of the fitness of its symbiotic partner (mediated through growth, feeding, and reproductive advantages)^7,8^. The benefits that the host incurs from their symbiotic partner may also vary with the species of the partner, e.g. specialized nutrients and protection^9–11^. However, as disturbances are intensifying and occurring more frequently, some host species are being disproportionally affected than other hosts^9,10,12^. In response, symbiotic partners may leave their host if it becomes unhealthy^11,13^, or they may stay and facilitate their mutual recovery^6,10,14^.

On coral reefs, corals are host to many mutually symbiotic organisms, such as microbes, *Symbiodinium* algae, crabs and coral-dwelling fishes^15–17^. These symbiotic partners often specialize on particular host coral species, which they may leave or stay during environmental stress^12,15–17^. Little is known about how climate change affects these mutualisms and the degree of host specialization by symbiotic partners, despite the importance of these ecological partnerships. For example, coral-fish symbioses are important for coral health because fish protect corals from toxic algae, sedimentation, predation, and stagnant hot water build-up^14,18–20^. Often, coral-dwelling fishes specialize on different hosts and vary to what extent they are specialized: some only live in 1-3 species (host specialist), while others use 4-11 coral species (host generalist)^7,12,21^. Host specialization by coral-dwelling fishes likely affects how both symbiotic partners recover given that climatic disturbances affect some hosts more than others^22,23^.

Here, our 7-year study (2013-2020) shows that coral-dwelling gobies (genus *Gobiodon*) either disappeared or shifted their occupation of host corals (genus *Acropora*) after an unprecedented succession of disturbances with limited recovery periods: 2 category 4 cyclones (2014, 2015) and 2 prolonged heatwaves (2016, 2017) which caused extensive coral bleaching. By surveying gobies and their coral hosts before and after each disturbance, and then 3 years post-disturbances, we found that gobies fared far worse than corals, with a distinct time lag in the early signs of recovery of gobies compared to corals^**23**^. Previous studies have shown trade-offs between goby fitness and host specificity, with particular coral hosts improving growth and survival of specialist gobies compared to generalist gobies^7,21^. Accordingly, the shifts in host occupation coupled with a lag in recovery of gobies will likely hamper fitness of both parties during the crucial and early stages following disturbances^14,18–20,24^.

## Results

### Goby Recovery is Lagging Behind the Recovery of their Coral Hosts

Throughout these consecutive disturbances and 3 years post-disturbances, we surveyed 36 species of *Acropora* coral hosts used by 13 species of coral-dwelling gobies (*Gobiodon*) known to occur at Lizard Island, Great Barrier Reef, Australia (−14.687264, 145.447039, Suppl. Fig 1, Fig 1A). Less than one year after the last disturbance (2018), coral and goby abundances, richness, coral diameter and occupancy, and goby group size were at an all-time low (Suppl. Fig 2, p < 0.001, see Supplementary Table 1 for all statistical results). Three years post-disturbances (2020), there were signs of recovery for corals as coral abundance and richness were higher than previously recorded, but coral diameter remained extremely small and corals were rarely occupied by gobies (Suppl. Fig 2). Goby richness and abundances were still very low, and gobies continued to occur singly (Suppl. Fig 2). However, the number of juvenile goby species and their abundance improved (Suppl. Fig 2).

**Fig 1.**
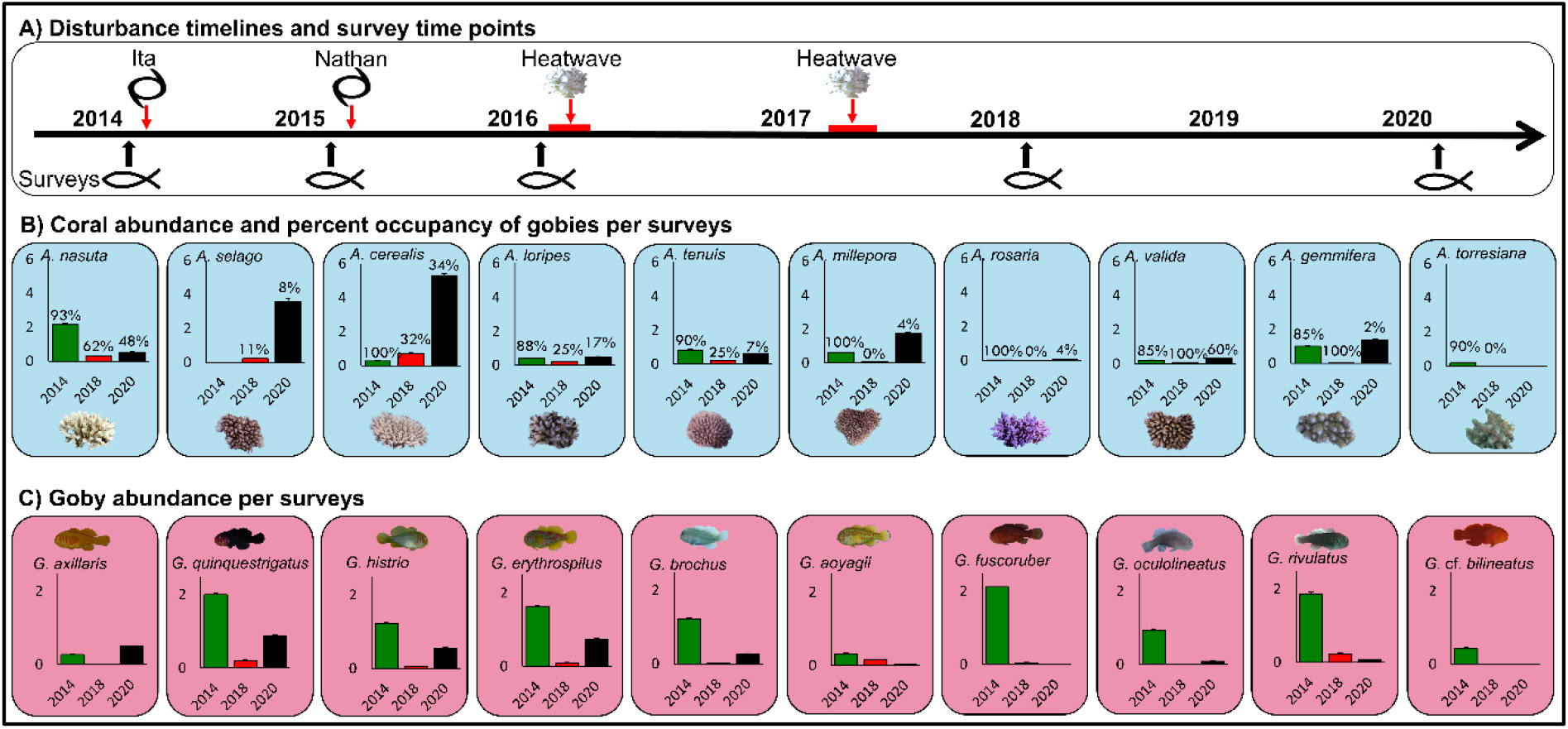
Multiple disturbances changed the mean abundance per transect of *Acropora* corals (blue) and their symbiotic *Gobiodon* gobies (red). A) Following consecutive disturbances (2 cyclones and 2 heatwaves), B) the 10 most common coral hosts and C) their goby symbionts experienced drastic changes in abundances. Abundances after each cyclone were not significant but were significant after the last disturbances, and thus we display changes post-disturbances. Error bars are standard error. Percentages above bars represent the proportion of corals that were occupied by gobies during that particular survey year.

We focused specifically on the abundance of the 10 most commonly used coral hosts and 10 most common goby species, and found that not all goby and coral species responded in the same way. Abundances were different among years (p < 0.001, Fig 1B), with eight coral species becoming extremely rare after disturbances, which was not surprising because 50% of the transects lacked corals compared to only 5% before disturbances^23^. However there was recovery 3 years post-disturbances when only 17% of transects lacked corals. Surprisingly, two coral species became more abundant immediately after disturbances even though they were rare before (*A. cerealis* and *A. selago*). These species became at least 10 times more abundant 3 years post-disturbances than pre-disturbances (Fig 1B). In general, more corals were found without goby partners post-compared to pre-disturbances (Fig 1B).

For gobies though, it was a different story. Several species were still absent three years post-disturbances (2020) (Fig 1C). Three species disappeared altogether from our survey sites immediately after disturbances (*G*. cf. *bilineatus, G. fuscoruber*, and *G. oculolineatus*), and an additional two species (*G. aoyagii*, and *G. rivulatus*) disappeared 3 years post-disturbances (Fig 1C). Of those species that disappeared, three were already rare before disturbances, but one was originally the most common species surveyed (*G. fuscoruber*). Only one goby (*G. axillaris*) returned to its pre-disturbance abundance in 2020 i.e. had fully recovered, while the remaining half that were still observed were still at 50% pre-disturbance abundances (Fig 1C).

### Some Gobies Showed Plasticity in their Host Specificity

Pre-disturbances, each goby species usually inhabited a range of coral species with minimal overlap among goby species (p < 0.01), but this variation in host specificity was affected by the climatic disturbances (p < 0.01, Suppl. Fig 3, Fig 2). Not all gobies responded the same in terms of host occupation throughout the disturbances (p < 0.01; Fig 2), although there were no marked differences in particular coral species occupied by host specialists versus host generalists (p > 0.50; Fig 2).

**Fig 2.**
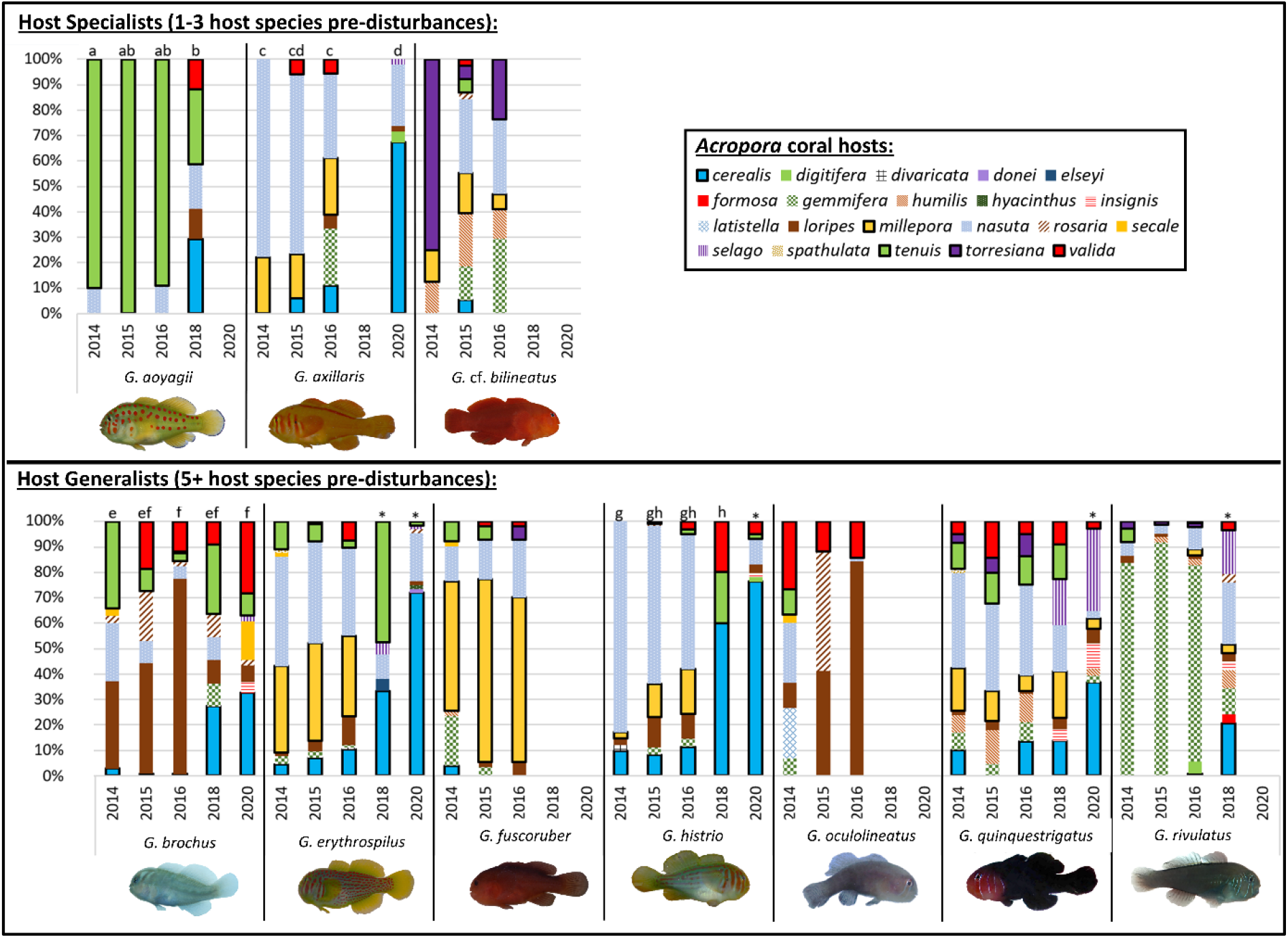
Host specificity of *Gobiodon* gobies in *Acropora* coral hosts changed following multiple disturbances. Proportion of all *Acropora* species used by the 10 most common *Gobiodon* species from surveys: pre-disturbances (2014), after cyclone Ita (2015), after cyclone Nathan (2016), after two back-to-back heatwaves/bleaching events (2018), and 3 years post-disturbances (2020). Letters above each bar represent host use differences among sampling years that are significantly similar to one another within species, and asterisks represent host occupation that is significantly different from all others within a species.

Host specialists, i.e. *G. aoyagii, G. axillaris*, and *G. cf. bilineatus*, occupied 1-3 host species pre-disturbances but each species occupied their own range of host species (Fig 2). Cyclones had minimal effects on host occupation, but there were marked changes after heatwaves. Post-disturbances, host specialists either disappeared or occupied more host species than previously observed (Fig 2). Of the three host specialists, *G*. aoyagii was the only species that was present after disturbances (2018) but it switched to being a host generalist occupying 5 coral species. Three years post-disturbances, G. ao*yagii* disappeared, but *G. axillaris* was observed once again and was a generalist occupying 5 coral species.

The other seven goby species were host generalists inhabiting between 5 to 10 coral host species pre-disturbances (Fig 2). Cyclones had minimal effect on host occupation, but heatwaves again caused noticeable changes. Post-disturbances, out of the seven host generalists, 5 goby species were still present and all but *G. histrio* remained host generalists, although *G. histrio* was only observed 10 times (Fig 2). Even three-years post disturbances, generalists continued occupying a wide range of hosts, including *G. histrio* again, although another generalist *G. rivulatus* had disappeared (Fig 2).

To index host specificity along a continuum instead of finite categories (host specialist vs. generalist), we calculated the proportion of occurrences that a goby species only occupied one coral species. We found that this index affected the range of hosts occupied throughout disturbances (p < 0.01); i.e., goby species that tended to occupy only one coral species occupied different coral species to goby species that tended to occupy several coral species. However, each goby species preferred a single particular coral species over others (Fig 3). In particular, gobies occupied a particular host between 35-90% of the time, although host specialists tended to occupy one host species more often than host generalists. For host specialists, 90% of *G. aoyagii* occupied *A. tenuis*, 75% of *G. axillaris* occupied *A. nasuta*, and 75% *G*. cf. *bilineatus* occupied *A. torresiana* (Fig 2,3). For host generalists, 30% of *G. brochus* occupied *A. loripes* and 30% occupied *A. tenuis*, 40% of *G. erythrospilus* occupied *A. nasuta*, 50% of *G. fuscoruber* occupied *A. millepora*, 80% of *G. histrio* occupied *A. nasuta*, 25% of *G. oculolineatus* occupied *A. valida*, 35% of *G. quinquestrigatus* occupied *A. nasuta*, and 80% of *G. rivulatus* occupied *A. gemmifera* (Fig 2,3). Therefore, *A. nasuta* was the preferred host for four goby species, whether they were host specialists or generalists (Fig 3).

**Fig 3.**
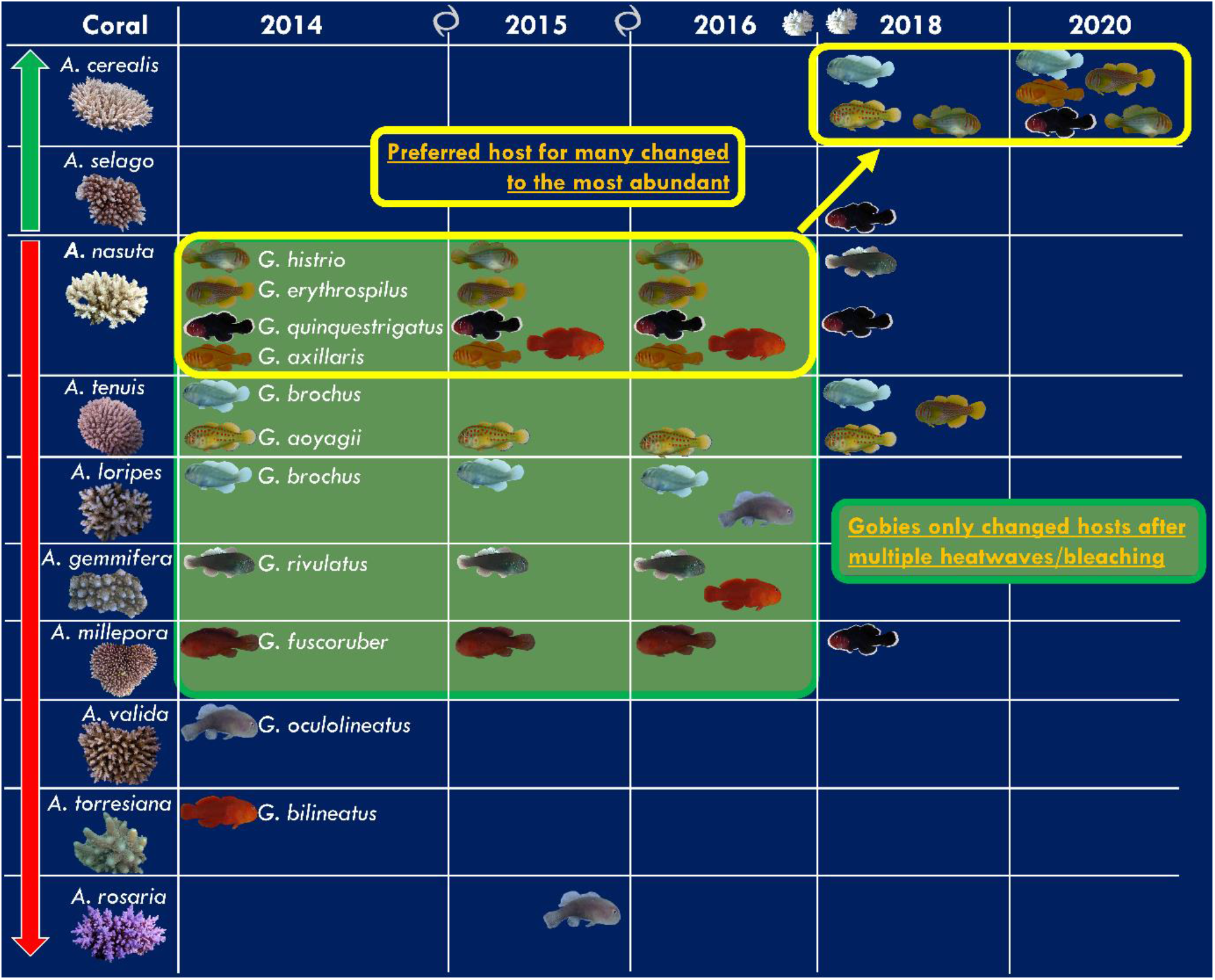
Changes in preferred *Acropora* host(s) for each *Gobiodon* gobies following multiple disturbances. Completed surveys before disturbance (2014), after cyclone Ita (2015), after cyclone Nathan (2016), after two back-to-back heatwaves (2018), and 3 years post-disturbances (2020). Coral hosts are organized from top to bottom to illustrate changes from most abundant to least abundant corals after disturbances. Green arrow highlights coral species that increased in abundance after disturbances, and red arrow highlights coral species that decreased in abundance after disturbances. Green box signifies gobies that did not change their preferred host until after heatwaves.

After the two cyclones, there was little change in preferred host, suggesting that cyclones did not alter host specificity^23^ (Fig 3). However after heatwaves, gobies shifted their host use, and often this shift mirrored the change in coral community. Many gobies switched from the previously popular *A. nasuta* to the newly abundant *A. cerealis* (Fig 1,3). Out of the remaining goby species post-disturbances, *Gobiodon* aoyagii began occupying *A. tenuis* and *A. cerealis* each 25% of the time, *G. histrio* switched to occupying the newly abundant *A. cerealis* 60% of the time, and three others (*G. brochus, G. erythrospilus*, and *G. rivulatus*) were also found more often in *A. cerealis* than previously observed (at least 20% of the time). The occupation of any particular host coral was not above 45% for any goby species after heatwaves, except for *G. histrio*.

Three years post-disturbances, there was little change in the number of hosts occupied by each goby species, but the majority of gobies were primarily occupying *A. cerealis* as it was the most abundant (Fig 1,3). *Gobiodon axillaris* was observed once again but switched host to *A. cerealis* 65% of the time (Fig 2,3). For *G. histrio* and *G. erythrospilus*, their preferred host was *A. cerealis* (75, 70% respectively), others like *G. brochus* used *A. cerealis* albeit to a lesser extent (30%), and *G. quinquestrigatus* used *A. cerealis* (35%) and *A. selago* (30%). Accordingly, even three years post-disturbances, most gobies used *A. cerealis* over other coral species (Fig 3).

## Discussion

As multiple disturbances are becoming the norm, we find that mutualisms on coral reefs may not respond as a collective unit. Our 7-year study shows that *Acropora* corals are faring better than their goby inhabitants (genus *Gobiodon*) 3 years after back-to-back climatic events (2 cyclones and 2 heatwaves)^23^. Right after disturbances, populations of corals and gobies were each devastated, but gobies declined at least three times more than corals and most corals were devoid of gobies^23^. After 3 years of recovery time, coral hosts became twice as abundant and speciose compared to pre-disturbances, although coral sizes were three times smaller than pre-disturbances. Reduced competition for space among corals may have allowed a surge in abundances within a few years of recovery, although they also had to compete with fast growing algae and high incidences of corallivory^24,25^. For gobies though, half of the goby species became rare or absent 3-years post-disturbances, and there were four times fewer adult gobies compared to pre-disturbances. In addition, these gobies were living singly, which suggested low turnover rates since gobies need to live in pairs to reproduce^26^. Since corals remained very small, gobies may have been unable to pair and breed as they need larger corals to do so^27^. As such, gobies may be facing a population bottleneck^28^ due to the inability to form pairs over multiple years. Alarmingly, 75% of corals no longer hosted gobies post-disturbances compared to just 5% pre-disturbances^23^. Even with 3 years of recovery time, 75% of corals were still devoid of gobies. Such a lag in goby resilience is dire for the mutualism of corals and gobies.

Given that habitat specificity is likely to play a key role in the continued prevalence of coral and goby symbioses, the fact that gobies shifted their host occupancy is a cause for concern. Initially, one third of the *Gobiodon* species inhabited just 2-3 host species, while others occupied a broader range of hosts ^16,29^. The disappearance of half of the goby species mirrored the decline in their preferred coral hosts immediately after cyclones and heatwaves. Thus, despite being an advantage during stable periods, being a host specialist may be a significant disadvantage during unstable periods, like those being fraught with multiple environmental disturbances^29,30^. Even more alarmingly, these specialist species stayed rare or disappeared despite their preferred host species increasing in abundance 3 years post-disturbances. Although these unoccupied corals may be able to survive in the short term, a prolonged lack of mutualistic goby partners may increase their vulnerability to external threats in the long-term since gobies provide beneficial services to corals^14,18–20,24^. However, it is possible that other goby species may shift hosts in the short-term, given the host plasticity observed in some species. Such host shifts may increase coral resilience but potentially decrease goby fitness, since goby growth rates are higher in their preferred coral species^7,21^. Over the longer-term, while the capacity for host shifts may promote initial short-term survival of both partners, the fitness of both gobies and corals may decline over time unless other coral symbionts fill the symbiont niche^12,15,17^. Given that disturbances are occurring more frequently than ever before^1,2^, the mutualism between coral hosts and gobies may not be able to persist after continued disturbances, leaving both organisms susceptible to additional stress that could even have knock-on effects on ecosystem stability^1,10,31,32^. Our study is an early warning sign that mutually symbiotic partners may not recover at similar rates, and while the capacity for plasticity in host occupation may be key for immediate survival, it may not prove a sustainable strategy for resilience to future environmental and other stressors.

## Methods

### Study Location

All sampling was completed at reef sites within Lizard Island, Great Barrier Reef, Queensland, Australia (−14.687264, 145.447039, Suppl. Fig 1). Lizard Island was affected by four extreme climatic events annually from 2014 to 2017: cyclone Ita (category 4) in April 2014, cyclone Nathan (category 4) in March 2015, heatwave causing a mass-bleaching event from March to April 2016, and a second heatwave causing a mass-bleaching event from February to May 2017. Sites were visited before these events in February 2014 (n = 18 sites), after the first cyclone in January-February 2015 (n = 16), after the second cyclone in January-February 2016 (n = 19), after both heatwaves in February-March 2018 (n = 22), and 3 years after the last disturbance in January-March 2020 (n = 24) (Fig 1A). Not all sites were sampled each year due to weather conditions and scouring effects of cyclones that left some sites with only bare rock.

### Sampling Method

Surveys were completed at each time point for the presence of *Gobiodon* goby spp. within *Acropora* coral spp. There were two types of surveys used: (1) in 2014, 2018, and 2020, corals were surveyed 1 m on either side of 30-m transects, and (2) in 2015 and 2016, corals were surveyed 1 m on either side of 4-m cross-transects^23,33^. In addition, since very few corals were encountered along transects after the four disturbances, random searches occurred in 2018 and 2020. When a live *Acropora* coral was encountered, the coral was measured and averaged along its width, length, and height^34^. Only corals at least 7 cm in average diameter were included in surveys, because smaller corals were never found occupied by gobies^23^. The coral was searched for a *Gobiodon* species using a bright torch light (Bigblue AL1200NP), and the species and number of individuals were noted. Individuals were identified as adults or juveniles based on coloration and size. The study was completed under the animal ethics protocols AE1404 and AE1725 from the University of Wollongong, and research permits G13/36197.1, G15/37533.1 and G18/41020.1 issued by the Great Barrier Reef Marine Park Authority.

### Data analysis

For changes in coral and goby populations, we used data from transects only since random searches did not follow any particular transect techniques. The following variables had many zero data points per transect after multiple disturbances, and accordingly were compared among survey yr (fixed factor) and site (random factor) with a generalized linear mixed model (GLMER: poisson family) using a zero-inflated model: coral richness and abundance, adult goby richness and abundance, and juvenile goby richness and abundance. Note: for all abundance variables, only line transects in 2014, 2018, and 2020 were used to remove transect type bias in abundances. The following variables were compared among survey yr (fixed factor) and site (random factor) with linear mixed models (LMER): average coral diameter, coral occupancy (whether occupied or unoccupied by *Gobiodon* spp.), and adult goby group size (juveniles were not included because they were observed moving between coral heads). All analyses were completed in R (v3.5.2)^35^ with the following packages: tidyverse^36^, lme4^37^, lmerTest^38^, LMERConvenienceFunctions^39^, piecewiseSEM^40^, glmmTMB^41^, emmeans^42^, DHARMa^43^, and performance^44^. Coral and goby communities for the 10 most common species of each genus were compared among survey yr (fixed factor) and site (random factor) with a permutational analysis of variance (PERMANOVA) in Primer-E software (v7).

For host specificity analyses, we used data from transects and random searches. Data for particular species were removed for years in which the species was observed less than 8 times in order to allow for enough observations to assess host specificity use. Three out of the 13 goby species observed in the surveys were excluded for host specificity analysis since they were consistently too rare (*G. citrinus, G. okinawae*, and *G. sp. D*). The corals inhabited per goby species were then combined within current zones per year. Coral species inhabited were compared among goby species (fixed factor) and survey yr (fixed factor) using PERMANOVA. The following covariable was added to the analysis which was calculated from the first survey pre-disturbances (2014): specificity continuum (proportion of occurrences in which only one coral species was used per goby species [continuous variable, 0-1]). PERMANOVAs were repeated (without the covariable as it is correlated with the following factors) to individually include each of the following explanatory factors calculated from the first survey pre-disturbances (2014): coral richness specificity (fixed factor, host specificity category per goby species on the basis that goby conspecifics used up to 3 coral species [specialist] versus more than 3 coral species [generalist]), proportional coral specificity (fixed factor, host specificity category per goby species on the basis that 75% or more goby conspecifics used a single coral species [specialist] versus less than 75% of gobies used a single coral species [generalist]), and sociality index of each goby species (fixed factor: asocial or social as calculated in Hing et al., 2018). Note: the goby species factor was nested within each of the factors in the later PERMANOVAs.

