## Supplementary Data for "Delayed recovery and host specialization may spell disaster for coral-fish mutualism"

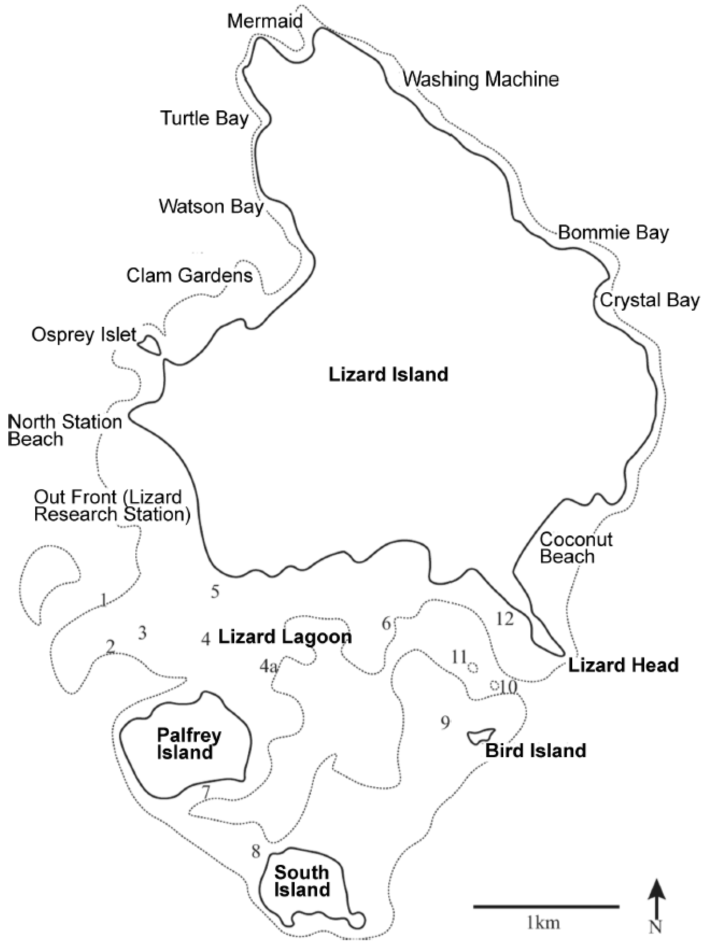


**Supplementary Fig 1. Map of Lizard Island, QLD (-14.687264, 145.447039).** Numbers represent discrete sites as follows: Big Vickey's Reef (1); Vickey's Reef (2); Horseshoe Reef (3); Palfrey Reef (4±4a); Loomis Reef (5); Trawler (6); Picnic Beach (7); Ghost Beach (8); Bird Island Reef (9); Entrance Bommie (10); Bird Bommie (11); Coconut Head Reef (12).

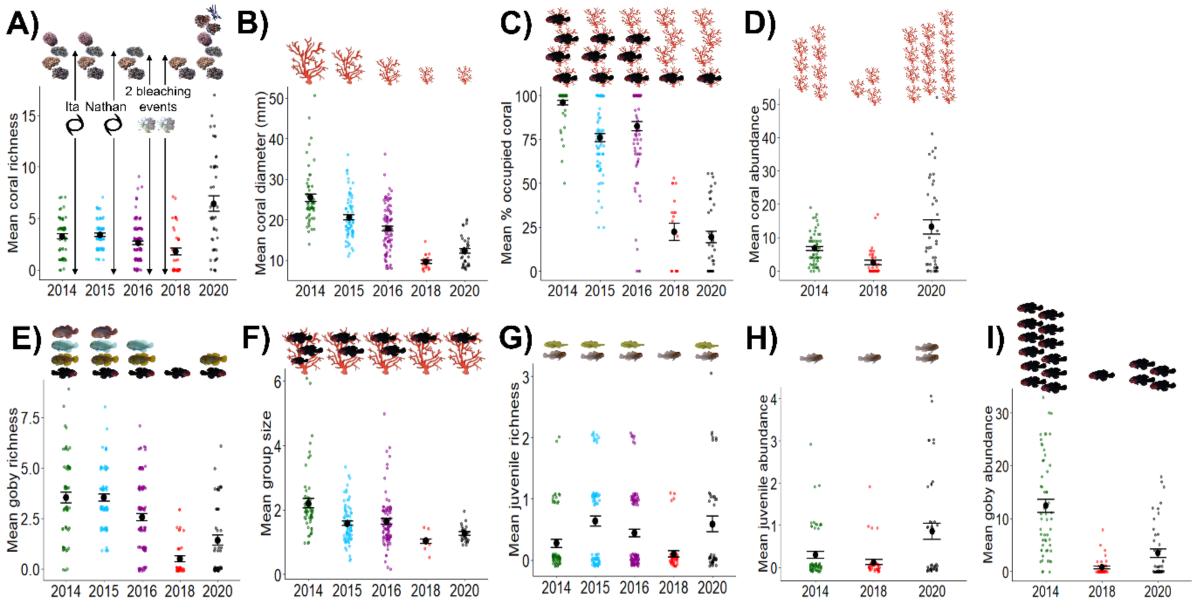


**Supplementary Fig 2. Multiple disturbances affect the populations of *Acropora* corals and *Gobiodon* gobies.** (A) Coral richness, (B) diameter,(C) occupancy by gobies, and (D) abundances, (E) adult goby richness and (F) group size, (G) juvenile goby richness, and the abundance of (H) juvenile gobies, and (I) adult gobies per transect before and after each cyclone (black spiral symbol), after heatwaves/bleaching (white corals), and 3 yr post-disturbances. Abundances (D,H,I) were only calculated in 2014, 2018, and 2020 due to differences in transect methodology. Symbols above data points depict changes in means as identified through post-hoc testing. Error bars are standard error.

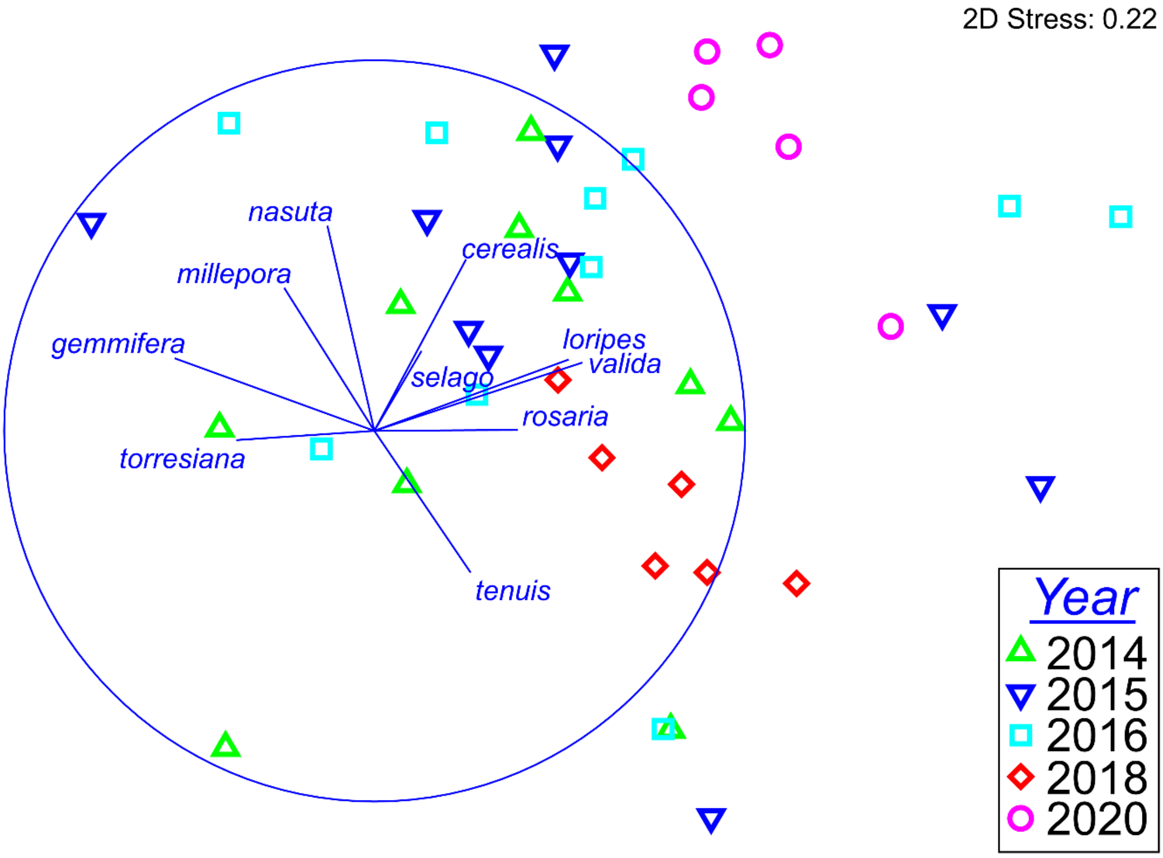


**Supplementary Fig 3. Coral assemblages inhabited by each goby species following multiple disturbances.** Multidimensional scaling plot illustrating surveys were completed: before (2014), after Cyclone Ita (2015), after cyclone Nathan (2016), after two back-to-back bleaching events (2018), and 3 yr after disturbance (2020). Each point represents the diversity of corals used by each goby species within a single survey time period. Points closer together are more similar in coral assemblages than points further apart. Overlayed are the direction in which *Acropora* coral species are most abundant, and the closer the trajectory is to reaching the outer circle, the most abundance of that species is explained in that direction.

Supplementary Table 1.Output of all statistical analyses. GLMM = generalized linear mixed model, LMM = linear mixed model, PERMANOVA = permutational analysis of variance

| Response Variable | Model | Predictor variable | Factor Type | df | Model Statistic | Test-value | p-value | R-squared |
|---|---|---|---|---|---|---|---|---|
| Coral Richness | GLMM | Year | fixed | 3 | $\chi^2$ (chi- | 87.395 | < 0.0001* | 0.185 |
| | poisson | Site | random | | squared) | | | marginal |
| | zero-inflated | | | | | | | |
| | | | | | Sample size | n = 326 | | |
| | | | | | Outliers removed | n = 9 | | |
| Average Coral Diameter | LMM | Year | fixed | 3 | F-value | 64.158 | < 0.0001* | 0.562 |
| log-transformed | | Site | random | | | | | conditional |
| | | | | | Sample size | n = 283 | | |
| | | | | | Outliers removed | n = 4 | | |
| Percent Occupied | LMM | Year | fixed | 4 | F-value | 132.21 | < 0.0001* | 0.723 |
| Corals | | Site | random | | | | | conditional |
| | | | | | Sample size | n = 283 | | |
| | | | | | Outliers removed | n = 6 | | |
| Adult Goby Richness | GLMM | Year | fixed | 7 | $\chi^2$ (chi- | 115.35 | < 0.0001* | 0.5 |
| | poisson | Site | random | | squared) | | | marginal |
| | zero-inflated | | | | | | | |
| | | | | | Sample size | n = 326 | | |
| | | | | | Outliers removed | n = 4 | | |
| Average Goby | LMM | Year | fixed | 4 | F-value | 10.471 | < 0.0001* | 0.194 |
| Group Size | | Site | random | | | | | conditional |
| log-transformed | | | | | | | | |
| | | | | | Sample size | n = 283 | | |
| | | | | | Outliers removed | n = 3 | | |
| Juvenile Goby Richness | GLMM | Year | fixed | 7 | $\chi^2$ (chi- | 25.843 | < 0.0001* | 0.21 |
| | poisson | Site | random | | squared) | | | marginal |
| | zero-inflated | | | | | | | |
| | | | | | Sample size | n = 326 | | |
| | | | | | Outliers removed | n = 11 | | |
| Coral Abundance | GLMM | Year | fixed | 3 | $\chi^2$ (chi- | 259.68 | < 0.0001* | 0.306 |
| | poisson | Site | random | | squared) | | | marginal |
| | zero-inflated | | | | | | | |
| | | | | | Sample size | n = 146 | | |
| | | | | | Outliers removed | n = 5 | | |
| Juvenile Goby Abundance | GLMM | Year | fixed | 5 | $\chi^2$ (chi- | 18.898 | < 0.0001* | 0.371 |
| | poisson | Site | random | | squared) | | | marginal |
| | zero-inflated | | | | | | | |
| | | | | | Sample size | n = 146 | | |
| | | | | | Outliers removed | n = 5 | | |
| Adult Goby Abundance | GLMM | Year | fixed | 3 | $\chi^2$ (chi- | 198.36 | < 0.0001* | 0.674 |
| | poisson | Site | random | | squared) | | | marginal |
| | zero-inflated | | | | | | | |
| | | | | | Sample size | n = 146 | | |
| | | | | | Outliers removed | n = 5 | | |
| Common 10 coral community | PERMANOVA | Year | fixed | 2 | pseudo-F | 7.958 | 0.0001* | N/A |
| | | Site | random | 16 | | 3.401 | 0.0001* | |
| | | Year*Site | | 32 | | 1.874 | 0.0001* | |
| | | Residual | | 95 | | | | |
| | | | | | Sample size | n = 146 | | |
| Common 10 goby community | PERMANOVA | Year | fixed | 2 | pseudo-F | 10.963 | 0.0001* | N/A |
| | | Site | random | 17 | | 3.465 | 0.0001* | |
| | | Year*Site | | 31 | | 1.719 | 0.0001* | |
| | | Residual | | 95 | | | | |
| | | | | | Sample size | n = 146 | | |
| Corals Used by Gobies | PERMANOVA | GobySpp | fixed | 9 | pseudo-F | 5.815 | 0.0001* | N/A |
| | | Year | fixed | 4 | | 9.58 | 0.0001* | |
| | | GobySpp*Year | fixed | 27 | | 1.288 | 0.005* | |
| | | Residual | | 124 | | | | |
| | | | | | Sample size | n = 165 | | |
| Corals Used by Gobies | PERMANOVA | Specificity Continuum | covariable | 1 | pseudo-F | 5.964 | 0.0001* | N/A |
| | | Year | fixed | 4 | | 9.538 | 0.0001* | |
| | | GobySpp | fixed | 8 | | 5.817 | 0.0001* | |
| | | GobySpp*Year | | 27 | | 1.288 | 0.005* | |
| | | Residual | | 124 | | | | |
| | | | | | Sample size | n = 165 | | |
| Corals Used by Gobies | PERMANOVA | Coral Richness Specificity | fixed | 1 | pseudo-F | 0.915 | 0.5124 | N/A |
| | | Year | fixed | 4 | | 6.189 | 0.0001* | |
| | | GobySpp(Coral Rich Spec) | fixed | 8 | | 5.934 | 0.0001* | |
| | | Coral Rich Spec*Year | fixed | 4 | | 1.466 | 0.0646 | |
| | | GobySpp*Year | fixed | 23 | | 1.21 | 0.038* | |
| | | Residual | | 124 | | | | |
| | | | | | Sample size | n = 165 | | |
| Corals Used by Gobies | PERMANOVA | Proportional Coral Specificity | fixed | 1 | pseudo-F | 0.81 | 0.587 | N/A |
| | | Year | fixed | 4 | | 5.829 | 0.0001* | |
| | | GobySpp(Prop Coral Spec) | fixed | 8 | | 5.898 | 0.0001* | |
| | | PropCoral Spec*Year | fixed | 4 | | 1.176 | 0.2474 | |
| | | GobySpp*Year | fixed | 23 | | 1.252 | 0.0197* | |
| | | Residual | | 124 | | | | |
| | | | | | Sample size | n = 165 | | |

Supplementary Table 1.Output of all statistical analyses. GLMM = generalized linear mixed model, LMM = linear mixed model, PERMANOVA = permutational analysis of variance

| Response Variable | Model | Predictor variable | Factor Type | df | Model Statistic | Test-value | p-value | R-squared |
|---|---|---|---|---|---|---|---|---|
| Corals Used by Gobies | PERMANOVA | Sociality Index | fixed | 1 | pseudo-F | 2.295 | 0.0491* | N/A |
| | | Year | fixed | 4 | | 6.515 | 0.0001* | |
| | | GobySpp(Sociality) | fixed | 8 | | 5.217 | 0.0001* | |
| | | Sociality*Year | fixed | 3 | | 2.104 | 0.004* | |
| | | GobySpp*Year | fixed | 24 | | 1.138 | 0.1139 | |
| | | Residual | | 124 | | | | |
| | | | | | Sample size | n = 165 | | |
